# Distribution of members of the *Anopheles gambiae* complex in selected forested tourist areas of Cross River State, Nigeria

**DOI:** 10.1101/805085

**Authors:** OA Oduwole, AO Oduola, CM Oringanje, NS Nwachuku, MM Meremikwu, MF Useh, AAA Alaribe

## Abstract

**Background:** The *Anopheles gambiae* mosquitoes are the most abundant, efficient and widely distributed vectors of the malaria parasite in sub-Saharan Africa. In most African countries, where malaria control programmes are focused on the use of long-lasting insecticide treated bed net, there is need to evaluate the biting behaviour and the identity of such mosquitoes to determine the relevance and appropriateness of the control measure implemented.

**Method:** This study investigated the distribution and molecular characteristics of the *Anopheles* species in selected forested areas in Cross River State, Nigeria. Mosquitoes were collected using pyrethrum spray catch and Centre for Disease Control light traps modified with yeast and sugar to generate carbon dioxide (CO_2_). *Anopheles gambiae* complex was identified using multiplex polymerase chain reaction followed by restriction fragment length polymorphism (PCR-RFLP) for molecular forms characterization.

**Results:** One hundred and four *Anopheles gambiae* s.*l*. were collected during the study. Multiplex PCR showed 75% of the species complex were *A. gambiae s.s*. and further characterization using PCR-RFLP showed that 53.8% of the *A. gambiae s.l*. identified were *A. gambiae s.s*.while 24.4% were *A.coluzzii*. The two species of the *A. gambiae s.l*. were found to be most abundant. The study also reported a 1.3% hybrid form of *Anopheles gambiae s.s*.and *Anopheles coluzzii*.

**Conclusion:** The findings suggest the first documented evidence of hybrid forms of *A. gambiae s.s*./*A.coluzzii* in South Eastern Nigeria although its epidemiological implication is still not clear.

## Background

Although reports show a remarkable reduction in the prevalence of malaria in sub-Saharan Africa, millions of people are still at risk of the disease in this region [1-3]. In southern Nigeria, malaria is holoendemic and occurs throughout the year with the highest transmission occurring between April and October during the wet season [4].

The control of malaria in Nigeria revolves around an integrated process which emphasizes prompt accurate diagnosis and treatment and the use of anti-vector control measures. The utilization of Long-lasting Insecticide Treated Bed Nets (LLINs) is one of the strategies employed by the National Malaria Elimination Programme for vector control [5]. This strategy focuses mainly on mosquitoes that feed indoors neglecting the species that feed outdoors.

The major malaria vectors in Nigeria are the *Anopheles gambiae* complex where in *A. gambiae s.s* and *A. arabiensis* are the most dominant sibling species, also the *Anopheles funestus* group [6-7]. These are widely distributed across Nigeria, covering the mangrove and coastal areas of the south, Guinea savannah in the middle belt to the Sahel savannah of the northern part of the country. *Anopheles gambiae* and *A. arabiensis* prefer to breed in an environment that is sunlit and has shallow temporary pockets of fresh water such as, puddles, pools and hoof prints and water collected in car tyre tracks [8]. In Nigeria, *Anopheles gambiae s.s*. has been reported to have an affinity for human blood (anthropophagic) and rests indoors (endophilic) with sporozoite rates ranging from 0.2 to 11.8% [8].In contrast, *A. arabiensis* has sporozoites rates of 0 to 4.8% and has been confirmed to have a preference for animal blood, feeding on humans in the absence of animals and resting outdoors (exophilic) [8]. The range and relative abundance of *A. gambiae s.s*. and *A. arabiensis* appear to be strongly influenced by climatic factors, such as total annual rainfall [9]. *A. gambiae s.s*.is prevalent in forested zones in contrast to *A. arabiensis* which is predominant in several Sudan Sahel and northern Guinea savannahs [10-11]. Generally, *A.arabiensis* tends to predominate in arid savannas, whereas *A.gambiae* is the dominant species in humid forest zones [12]. Where the two appear in sympatry, large changes in species composition often occur with *A.arabiensis* predominating during the dry season and *A. gambiae* becoming more abundant during the rainy season [11].

*Anopheles gambiae s.s*. is divided into five chromosomal forms as a result of a paracentric inversion on chromosome two [13-14]. In addition to the chromosomal differentiation, *A. gambiae s.s* was differentiated molecularly based on the sequence differences in the “intergenic spacer” (IGS) of the rDNA that is on the ‘X’ chromosome into ‘S’ and ‘M’ molecular forms [15]. Few years ago, a study reported that *A. gambiae* ‘M’ form is another species and not a genetic variation of *A. gambiae s.s*.and is now widely accepted internationally [16]. Thus, while the ‘S’ form retains the name *A. gambiae s.s*., the ‘M’ form is now known as *A. coluzzii*. It has been reported that *A. gambiae s.s*.and *A. coluzzii* exists in sympatry in West Africa. Studies in Lagos, South Western Nigeria which lies in the forest ecological zone of Nigeria [17-18] and Kano, Northern Nigeria in Savannah ecological zone of Nigeria [19] showed that *A. coluzzii* is more abundant than *A. gambiae*. However, this is in contrast with the previous report from a wider surveillance which showed that the molecular S form (now known as *A. gambiae*) is predominant and has a wider distribution across Nigeria compared to *A. coluzzii* [7-20] Hybridization of the two species is rarely reported to occur, however,the hybrid form was recently reported in Nigeria by two studies from South Western Nigeria [21] and North Central Nigeria [22] respectively. The rainforest belt of Nigeria where Cross River State (CRS) is located has a fairly large population of very rare wildlife. The state shares a long border with Cameroon to form a protected ecological zone and it is recognized by United Nations Educational, Scientific and Cultural Organisation (UNESCO) as a world heritage centre. The forests host up to 16 species of primates. These include Chimpanzees, Drill monkeys, potty-nosed monkeys, Mangabey monkeys, Preuss’s Guenon and many others [23-25]. Cross River state has an estimated population of over three million eight hundred thousand people and is a major tourist destination in Nigeria because of its rare eco-tourism. Some of these forests are located in Akamkpa and Boki Local government areas of CRS. Villagers living close to the forests also hunt the animals for food and go to the forests for logs; in addition, forest rangers live in some part of the forests to protect the animals.

There is a paucity of information on the identity of members of the *Anopheles gambiae complex* in forested communities that border the wildlife sanctuaries in South Eastern Nigeria, which is usually in the print for malaria for unprotected immune and non-immune foreign tourists. In addition to the threat of zoonotic diseases to tourists and humans living in these forested communities, it is expedient to evaluate the current malaria vectors driving malaria distribution in forest communities that are conserved for endangered non-human primates and other wildlife in Cross River State.

## Materials and Methods

### Study area

This study was carried out in forested areas and border communities of Cross River State, South Eastern Nigeria. Cross River State has an estimated population of about 3,800,000 million people who are mostly farmers.

Mosquitoes were collected in the following locations; Bonchor, Drill Ranch in Afi Mountain Wildlife Sanctuary (Latitude 6.2999^0^ and Longitude 8.9977^0^, Altitude 300m), Cross River National Park (Latitude 5.2130°, Longitude 8.2548°, Altitude 143), Obung (Latitude 5.3458, Longitude 8.39445) Aking/Osunba(Latitude 5.4334 Longitude 8.6370 Altitude 182.6), Iko-Esai and Rhoko forest, (Latitude 4.9714 Longitude 8.3216 Altitude 135m)(Figure. 1), and Calabar municipal, (Latitude 4.977°, Longitude 8.334°, Altitude 135m). Calabar Municipal was the only urban setting in the study area, and was included because of the two wildlife sanctuaries (CERCOPAN and Pandrillus) located in the town. The average annual rainfall measurement during the study period was about 2,863.5mm (obtained from CERCOPAN and Pandrillus, Nigeria). Villages were selected if they were close to wildlife reserves.

**Figure 1:**
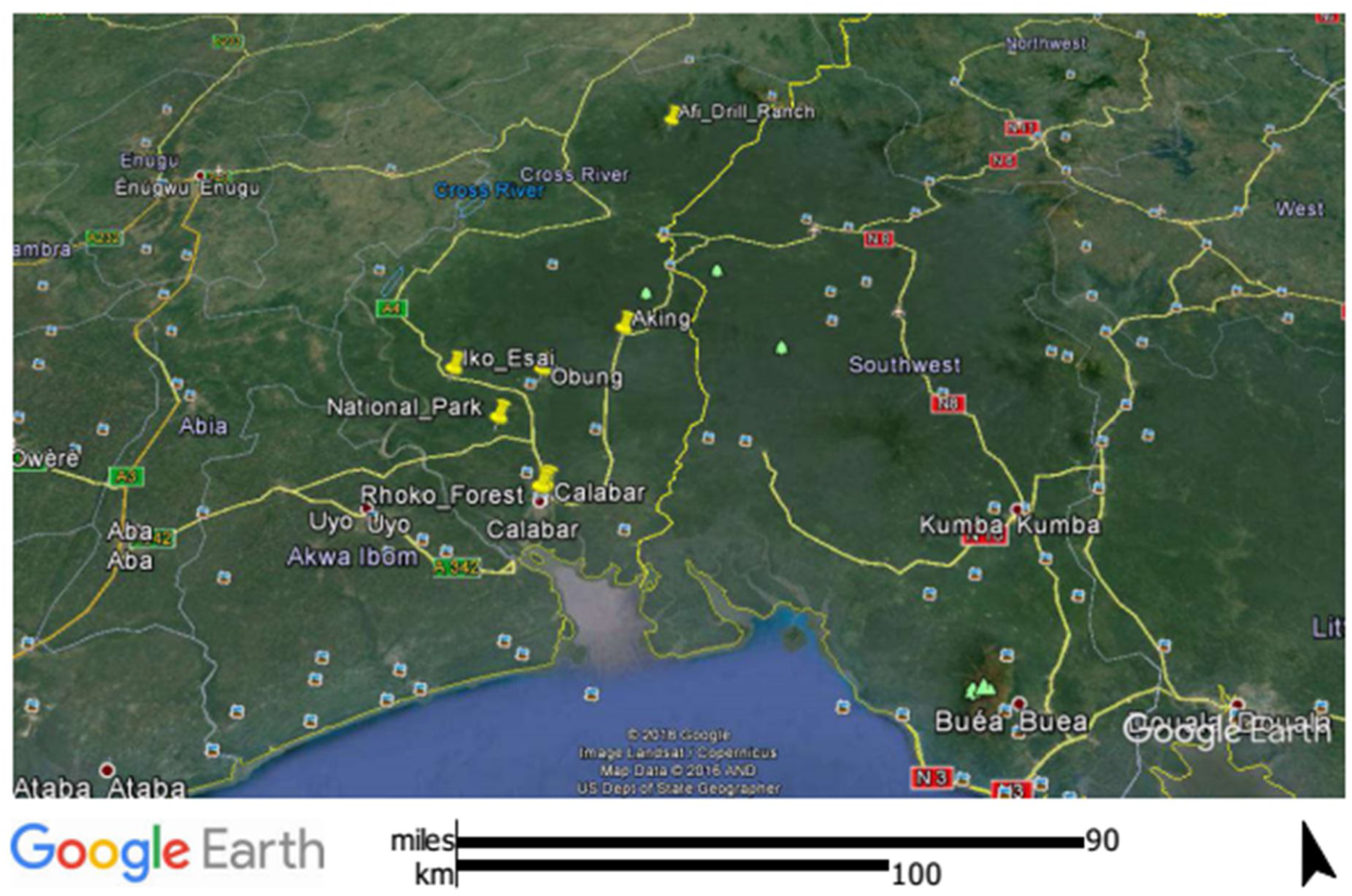
Geographic Positioning System (GPS) of Study Location Sample collection.

### Sample collection

Mosquitoes were collected using CDC light-traps and CDC ultraviolet light traps (Model 1312 and 912, manufactured by John W. Hock Company, Gainesville, Florida, U.S.A) modified with yeast and sugarto generate CO_2_ as described by Obenauer et al. [27] and synthetic lure (BG-Lure™ Biogents®, Regensburg). This was to enhance the attraction of the *Anopheles* to the light traps [27]. The CDC mosquito traps were set at different locations such as outdoors nearslow-flowing rivers or streams in the border communities and forests, outdoor close to human dwellings, and inside rooms where humans sleep under insecticide treated nets (ITNs). Six traps were set up near slow-flowing streams for three consecutive days in each community and six in the nearby forest. Communities with wildlife sanctuary were selected as forest locations. Each trap was about 15m from the other, following the method described in another study [27]. All traps were covered with a large black cover supplied with the traps to protect the trap from rain (Supplementary Figure 1). Six traps were also set up outside human dwellings about 15m away from living quarters, and in rooms where occupants sleep under ITNs. This is because the mosquitoes may not be attracted to the traps if they can bite humans. The mosquitoes were collected all night for three consecutive days monthly from 1800hours to 0600hours over a period of 12 months from April 2013 to June 2014 In addition, pyrethrum indoor spray catch was conducted from April 2013 to June 2014, in an average of 8 houses per study area perday between 0600hours and 0700hours for three consecutive days to cover dry (December to March) and rainy seasons (April to November). The knocked down mosquitoes were collected and kept in Eppendorf tubes in which silica gel had been added and plugged with cotton balls. Mosquitoes collected with the CDC traps were killed by keeping them in the -20°C freezer for two hours. For remote study locations without electricity, the traps were kept in bags already sprayed with insecticide. The female mosquitoes were later sorted out into different species and stored in an Eppendorf tube containing silica gel and plugged with cotton wool for further confirmation.

### Laboratory procedures

#### Morphological identification

*Anopheline* mosquitoes were separated from other genera of mosquitoes based on the characteristics of their maxillary palps. Also, males were differentiated from the females using their antennae, which are plumose for males and pilose for the females [28]. Furthermore, the female *Anopheles* mosquitoes were identified using well described morphological identification key [29-30]. After morphological identification, head and thoraces of the *Anopheles* were removed with a scalpel blade and examined for sporozoites using enzyme-linked immunosorbent assay technique as described by Obenauer et al. [27].

#### DNA Extraction

Deoxyribonucleic Acid (DNA) was extracted from individual *A. gambiae* placed in a 2ml Eppendorf tube using a QIAamp DNA mini kit (Qiagen). The individual mosquito was ground using plastic pestle. DNA extraction was completed following the manufacturer’s protocol. Eluted DNA was frozen at -20°C for further molecular analysis.

#### Mosquito species genotyping

Species identification was based on species-specific fixed differences in the rDNA region, including 28S coding region and intergenic spacer (IGS) region. DNA extracted from legs and wings of mosquitoes were subjected to species specific PCR assays following the procedure of Scott et al.[31]. Laboratory strains of the *Anopheline species* provided were used as positive controls. PCR products were visualized under UV light following gel electrophoresis. Positive amplicons for *A. gambiae s.s* were further digested to give rise to molecular forms S (formerly *A. gambiae)* and M (formerly *A. coluzzii*) using Restriction Fragment Length Polymorphism (RFLP) [32]. The result was analyzed by using 2% agarose gel electrophoresis and GelRed staining.

#### Statistical Analysis

Z-test, and Chi-square test were used to determine differences between *A. gambiae s.s.* (S form) and *A. coluzzii* (M form) and their association with variables such as season of the year, location (indoor or outdoor) and geographical area..

## Results

### Distribution of mosquito species in the study area

Data were obtained from the following study locations; Cross River National Park (CRNP), Obung, Osunba, Aking, Rhoko forest,and Iko-Esai in Akampka; Drill Ranch (Afi mountain wildlife sanctuary) and Bonchor in Boki and Calabar municipality. One thousand, one hundred and eighty-two (1182) mosquitoes comprising 10 known species from 7 genera were trapped between April 2013 and June 2014 using different trapping methods including pyrethrum indoor spray catch. Forty of these mosquitoes were caught using human bait catch. The measurement recorded in June 2013, was 403mm compared to 287mm recorded in June 2014 (Table 1). It was observed that fewer mosquitoes were collected during the heavy precipitation. In June 2013, the lowest distribution of 4% was recorded during the whole study as against the 23% collected in June 2014 which was the highest collection made during the period.

**Table 1:** Rainfall measurement during data collection and number of mosquitoes trapped

| Month of collection<br>(April 2013-June 2014) | Rainfall<br>measurement<br>(mm) | No. of Mosquitoes<br>Trapped (%) |
| --- | --- | --- |
| April | 181.1 | 23 (2%) |
| June | 405.1 | 46 (4%) |
| July | 507.8 | 12 (1%) |
| August | 218.5 | 15 (1%) |
| September | 544.5 | 46 (4%) |
| October | 450.6 | 112(10%) |
| December | 148.8 | 63 (5%) |
| January | 15 | 61(5%) |
| March | 81.1 | 19(2%) |
| April | 230.1 | 92(8%) |
| May | 349.6 | 418(35%) |
| June | 287.2 | 275(23%) |
| **Total** | **2563.8** | **1182(100%)** |

As expected, the number of mosquitoes collected in the wet season [1039 (88%)] was significantly higher than the mosquitoes collected in the dry season 143 (12%) (P<0.0002) Figure 2 shows a similar observation for the female mosquitoes collected for the study.

**Figure 2:**
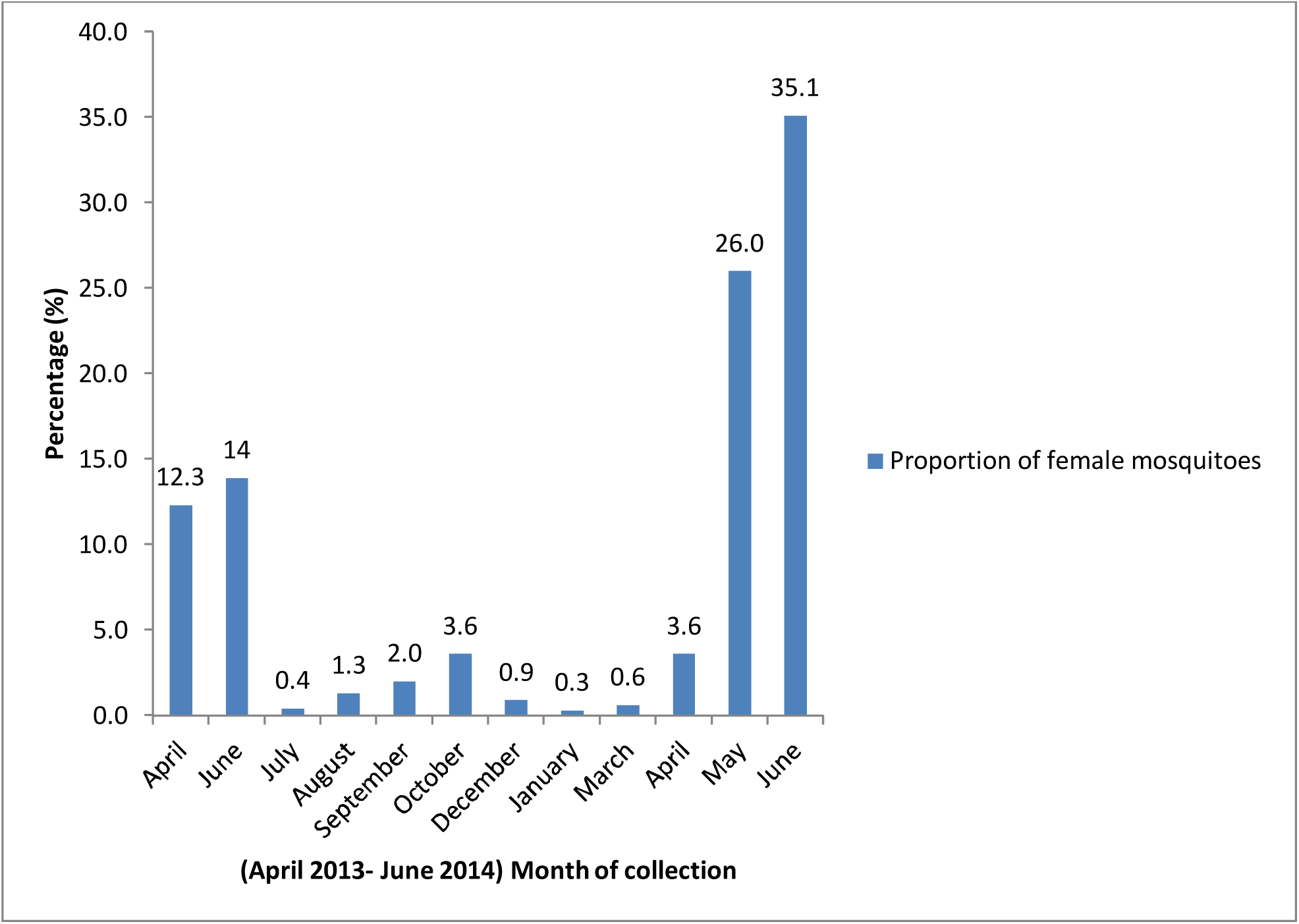
Distribution of female mosquitoes trapped by months of collection.

Seven hundred and seventy (770) of the mosquitoes were females, out of which 104 (13.5%) were *Anopheles species*. Other species were *Culex* species (54.4%), *Uranotaenia* species (13.3%), *Aedes* species (4.7%), *Mansonia* species (0.7%), *Lutzia species* (0.1%), *Coquillettidia* species (0.1%), while 13.1% could not be identified because they had become overgrown with fungi.*Culex* species were significantly more abundant than other species followed by *Anopheles* species (P< 0.0001). Rhoko forest had the highest proportion of female mosquitoes (22%), National park had19%, Calabar had18 %, Obung had 17% and Bonchor recorded the least 3% (Figure 3a)Obung had the highest proportion of female *Anopheles* mosquitoes (40%), followed by National park (30%) and Rhoko forest had the least proportion of female *Anopheles* mosquitoes(1.8%) (Figure 3b).

**Figure 3:**
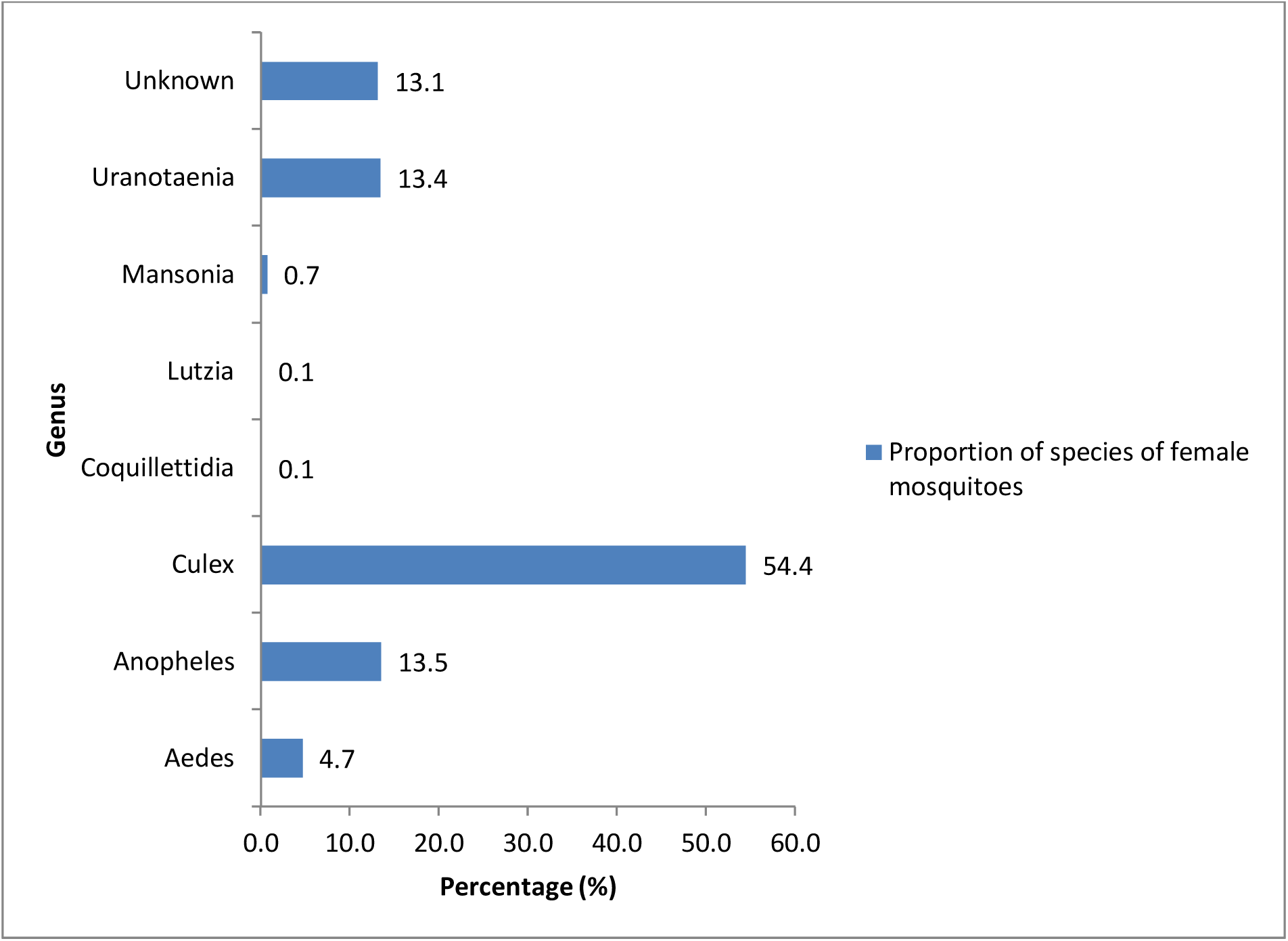
(a) Proportion of female mosquitoes caught.

**Figure 3:**
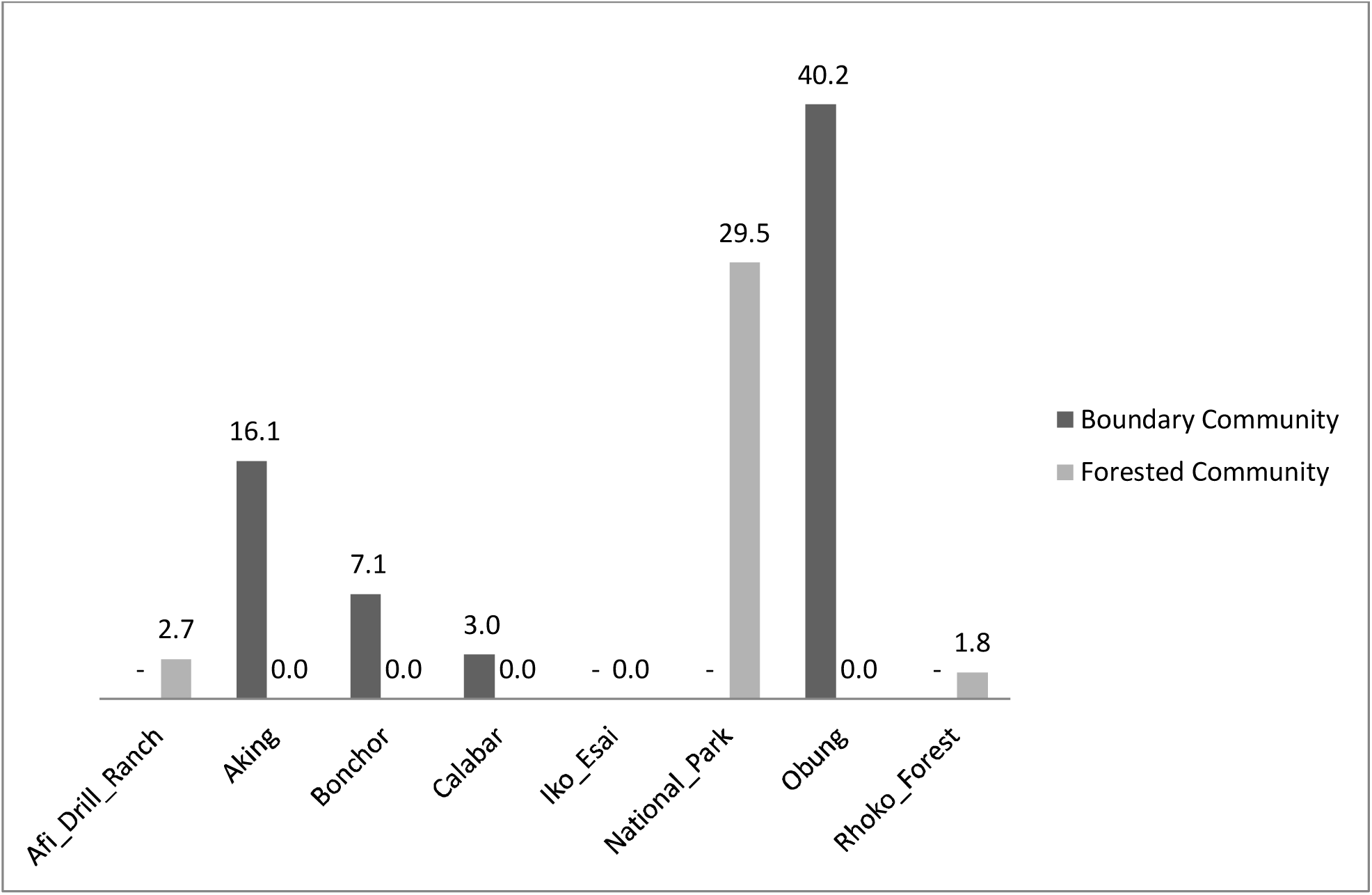
(b): Proportion of female *Anopheles* mosquitoes per study site.

Mophological identification shows that *A.gambiae* s.l. made up about 97% of the *Anopheles* species and 2.9% were *A. rufipes*. Among the identified *A. gambiae* s.l, 77% were *A. gambiae s.s* (P=0.0012); the remaining 23% could not be identified because of contamination caused by fungi (Supplementary Table 1).

*A gambiae s.s* samples were further classified into their molecular forms usingPCR-RFLP. Forty-two (53.8%) were identified as S form (*A. gambiae s.s*) and 19 (24.4%) were identified as M form (*A. coluzzii*) (Z=-6.1293, P-value<0.0001). Sixteen of the mosquitoes (20.5%) had no specific band and could not be classified into molecular forms, while 1(1.3%) was a hybrid form(S/M) (Figure 4).

**Figure 4:**
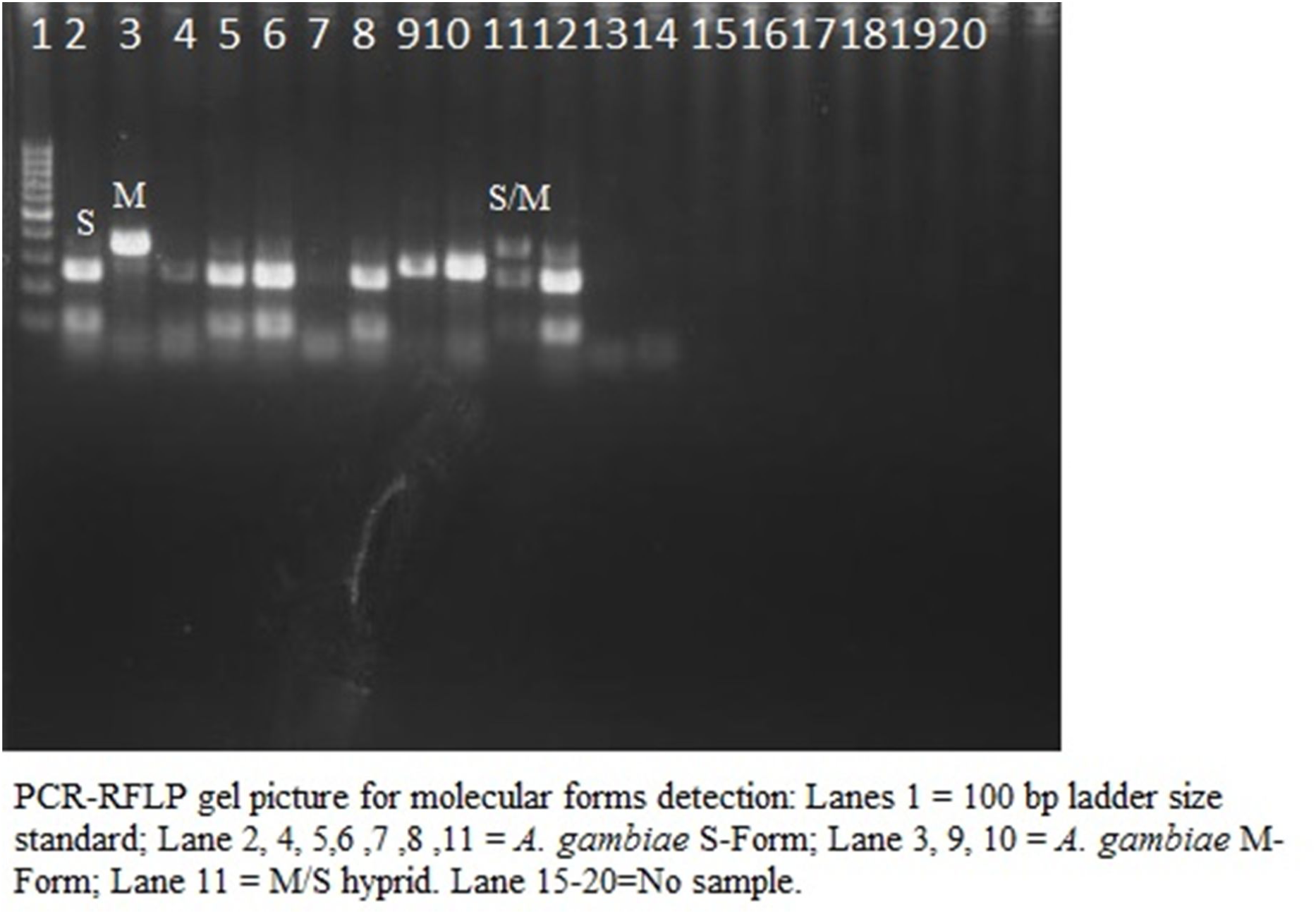
Molecular identification of *A.gambiae* s.s. and *A.coluzzii* using RFLP.

Overall, there was a higher distribution of *Anopheles* species in the border communities than the forests and more were caught indoors than outdoors of human dwellings within the communities. Also, for traps set outdoors, we collected more *Anopheles* species near slow-flowing streams than other locations outdoor. The mean number of the different genus of mosquitoes collected over the study period is represented in (Table 2). Analysis demonstrated a significant difference according to trap location. Mean distribution of *Anopheles* species, *Culex* species. (*p*<0.001) and other species (*p*=0.003) were significantly caught indoors than by the slow-flowing stream. Additionally, more *Anopheles* species (p<0.001), *Culex* species (*p*<0.001), *Aedes* species (*p*=0.002) and other species (*p*<0.001) were significantly caught outdoor of human dwellings than by the streams (Table 2). A similar capture trend was observed for the total mean number of mosquitoes irrespective of genus classification.

**Table 2:** Differences in the number (mean ± SD) of mosquito genus caught by trap location in 2013 – 2014 in selected forested tourist areas of Cross River State, Nigeria.

| Genus | Trap location |  |  |  |  |
| --- | --- | --- | --- | --- | --- |
|  | Stream | Indoor | Outdoor | F | p-value |
| Aedes (n=45)<br>(Row mean-Col mean), p-value | 1.40 ± 0.30 | 1.75 ± 0.03 | 2.11 ± 0.22 | 12.27 | 0.002* |
| Stream | - | - | - |  |  |
| Indoor | (0.35), 0.237 | - | - |  |  |
| Outdoor | (0.71), <u>0.002</u> | (0.36), 0.093 | - |  |  |
| Anopheles (n=104)<br>(Row mean-Col mean), p-value | 1.33 ± 0.34 | 1.77 ± 0.05 | 2.05 ± 0.15 | 26.78 | <0.001* |
| Stream | - | - | - |  |  |
| Indoor | (0.44), <u><0.001</u> | - | - |  |  |
| Outdoor | (0.72), <u><0.001</u> | (0.28), 0.056 | - |  |  |
| Culex (n=424)<br>(Row mean-Col mean), p-value | 1.31 ± 0.23 | 1.75 ± 0.06 | 2.02 ± 0.11 | 80.23 | <0.001* |
| Stream | - | - | - |  |  |
| Indoor | (0.44), <u><0.001</u> | - | - |  |  |
| Outdoor | (0.71), <u><0.001</u> | (0.26), <u>0.003</u> | - |  |  |
| Unknown (n=373)<br>(Row mean-Col mean), p-value | 1.27 ± 0.25 | 1.76 ± 0.06 | 2.07 ± 0.19 | 53.82 | <0.001* |
| Stream | - | - | - |  |  |
| Indoor | (0.49), <u><0.001</u> | - | - |  |  |
| Outdoor | (0.80), <u><0.001</u> | (0.31), <u>0.024</u> | - |  |  |
| **Other (n=175)<br>(Row mean-Col mean), p-value | 1.31 ± 0.23 | 1.75 ± 0.06 | 2.01 ± 0.12 | 53.42 | <0.001* |
| Stream | - | - | - |  |  |
| Indoor | (0.44), <u>0.003</u> | - | - |  |  |
| Outdoor | (0.70), <u><0.001</u> | (0.26), 0.171 | - |  |  |
| ***Total mosquitoes<br>(n=1,121)<br>(Row mean-Col mean), p-value | 1.30 ± 0.25 | 1.76 ± 0.06 | 2.05 ± 0.16 | 269.15 | <0.001* |
| Stream | - | - | - |  |  |
| Indoor | (0.46), <u><0.001</u> | - | - |  |  |
| Outdoor | (0.75), <u><0.001</u> | (0.29), <u><0.001</u> | - |  |  |

*#Lutzia genus was excluded in the analysis because of few observed number (only one mosquito was caught)*
*\*Significant p-value using one-way ANOVA test. Underlined p-values were significant after a post hoc Bonferroni test*
*\*\* Aedes, Mansonia, Uranotaenia species*
*\*\* \*Forty mosquitoes collected by human bait were not included in the analysis*

## Discussion

A higher collection of mosquitoes was observed in the wet season, during which one thousand, and thirty nine mosquitoes (88%) were collected, compared to 143 (12%) in the dry season. This observation was similar to findings from other study areas [33]. *Anopheles* abundance and malaria transmission is usually characterized and dependent on rainfall in Nigeria as this marks the availability of breeding sites [33]. This is probably because *Anopheles gambiae s.l*.is known to have a preference for clear water sources as their breeding grounds [28], which are readily available during the rainy season. The highest mosquito distribution was observed in June 2014 in contrast to the lowest in June 2013. This is may be due to the heavy rainfall recorded in June 2013, which was about twice of the measurement of rainfall in June 2014. It is believed that excess rainfall can wash away the mosquitoes’ breeding sites [34].

Morphological identification of the *Anopheles* species showed that *A. gambiae s.l*. and *A. rufipes* were the two *Anopheles* species identified. While *A. rufipes* is not a major vector of malaria, it has been implicated in some recent studies as a secondary vector of malaria [35]. This study also showed that *A. gambiae s.s* was significantly more abundant than *A.coluzzii*. This is similar to results from other parts of Nigeria [7, 20] showing that *A. gambiae s.s*.is a predominant and widely distributed species, especially in Southern Nigeria compared to the *A.coluzzii*.

In addition, this study found a hybrid form of *A. gambiaes.s./A.coluzzii*, which agrees with the findings from two recent studies [22]. The findings from our study and the previous reports showed that the hybridization of *A. gambiaes.s./A.coluzzii* is still rare with these two studies [21-22] reporting its occurrence in Nigeria at a low prevalence that ranges between 0.5% – 0.8 %. Although, the epidemiological implication of the hybrid form to malaria control is still unclear, it should be a cause for concern because of the possible transference of “knock down resistant gene” (kdr gene) from *A. gambiae* to *A.coluzzii* [32, 36-37].

As expected, more *Anopheles gambiae* species were found in the border communities compared to the forest locations but the difference was not significant. However, it was observed that a high proportion of the *A. gambiae s.s*. caught in this study were from the CRNP even though it is located in the forest. The CRNP has sleeping quarters built for rangers and tourists with wide open spaces which harbor pockets of water from rainfall and human activities. On the contrary, while tourists were allowed to visit the Drill Ranch and Rhoko forests, human activities that involved alteration of the forests in any form were discouraged. It is believed that small collections of water from rainfall and human activities will encourage *A. gambiaes.s*. to breed, in addition to the availability of blood meals from humans [6,21,38]. This was evident in the distribution of *A. gambiae s.s* in CRNP at 40% compared to 4% caught at Drill Ranch and none from Rhoko forest.

It was observed that in sites around human dwellings (outdoors and indoors), there were greater numbers of *Anopheles* mosquitoes collected than by the stream. This may be because most of the *Anopheles* species are *A.gambiae* and *A.coluzzi*, which are naturally attracted to human dwelling. Additionally, the *Anopheles* species collected outdoors were higher than the mosquitoes collected indoor but the difference was not statistically significant. It has been widely reported that *A. gambiae* is a species that is notorious for feeding and resting indoors [37]. The marginal decrease in the number of mosquitoes collected indoors compared to outdoors could be as a result of the malaria control programme, which is targeted only at indoor resting and biting mosquitoes resulting in a shift in behavior. There have been evidence on changes in biting and resting behavior of malaria vectors as a result of bed-net usage [39]. With 77% of the *Anopheles* species identified as *A.gambiae s.s*It is important to state that the presence of *Anopheles* species outdoor also has serious consequences for malaria control programme in the study area, since the main malaria control intervention involves the indoor use of LLINs. Tourists and residents enjoying outdoor evening time around this location may have to resort to the use of other alternatives such as repellents or wearing long clothing to protect themselves from mosquito bites. In addition, high levels of outdoor biting by *A. gambiae s.s*. have also been reported in a study in Equatorial Guinea [40]. This may corresponds to outdoor human activities in the early evening, however data on this was not collected.

This study did not detect the presence of *A. arabiensis*. This could be because *A. arabiensis* is predominantly found in arid environments and in areas where deforestation and urbanization have taken place [9,20]. In addition, there is a possibility of existence of *A. arabiensis* in the 25% unidentified *A.gambiae* complex. Furthermore, it is possible that the type of traps and where they were located could have played a major role in the number of mosquitoes caught during this study.

This study did not detect malaria infection in any of the malaria vectors identified. This may indicate to a certain extent that the malaria control strategies have been effective in creating a barrier between the mosquitoes and the hosts. The detection of infected *Anopheles* mosquitoes could also have been missed if the malaria parasites were still at the oocyst stage. This is in tandem with the observation in Oduwole et al. [26]. The report of the study showed that the prevalence of malaria among adults screened was low at 9.8% compared to the national average which is put at 27% by the WHO [41].

The high distribution of other species of mosquitoes such as *Culex* in large proportion should also be worrisome because of their ability to transmit neglected tropical diseases such as the lymphatic filariasis which is prevalent in Cross River State, Nigeria [42-45] In addition, *Aedes* species was about 5% of the mosquitoes collected during the study. This should be a source of concern because of its capacity to transmit yellow fever, a re-emerging viral haemorrhagic disease in Nigeria [3]. This study shows that tourists frequenting the forests and people living around non-human primates may not be at risk of simian malaria as observed in other studies [27, 46].

## Conclusion

*Anopheles gambiae* and *A. coluzzii* may be responsible for malaria transmission in the forest border between Cross River State and Cameroon but the existence of *A. rufipe* should not be taken for granted as secondary vector may constitute a nuisance in certain conditions. Additionally, hybridization of *A.gambiae* and *A. coluzzii* is still rare. Our study reported a prevalence of 1.3% that is not far from the range reported in previous studies from Nigeria and it is the first report from South Eastern Nigeria.

It is propose that tourists visiting these forests should wear protective clothing and use insect repellant during the day and sleep under LLIN at night to prevent malaria infection and other vector-borne diseases.

It is also recommended that future studies should investigate human behavior in the study area and how humans interact with the mosquito vectors.

## Study limitation

The main study limitation is that there were not many traps available per study site due to limited resources. This may have contributed to the low number of mosquitoes caught during the study.

## Authors’ contributions

OOA conceived the study. OOA, AAA and UMF designed the study. Laboratory investigation was supervised by OOA and NNS. Data collection was supervised by OOA and NNS and was interpreted by OOA, OAO, CO. OOA and MMM wrote the first draft of the manuscript. All authors criticallyappraised and approved the final draft of the manuscript.

## Compliance with ethical standards

Not applicable.

## Consent for publication

.Not applicable.

## Availability of data and materials

The data sets generated during the current study are available from the corresponding author on reasonable request.

## Abbreviations

(CRS): Cross River State
(PCR-RFLP): Restriction fragment length polymorphism
(WHO): World Health Organization
(LLIN): Long Lasting Insecticide Treated Bed Net
(IGS): intergenic spacer
(CRNP): Cross River National Park
(CDC): Centre for Disease Control
(CDC UV): Centre for Disease Control Ultraviolet
(Spp): Species
(DNA): Deoxyribonucleic Acid
(NAMRU - 3): Naval Medical Research Unit number -3

## Acknowledgements

This study was supported in part by theTertiary Education Trust Fund (TETFUND) and US-Naval Medical Research Unit Number-3 (NAMRU-3). We would like to thank Friday O. Odey for assisting with the field work, Iwara Arikpo and Okoro Anthony for data analysis, Nahmy Fahmy and Mba Mosore for assisting with the laboratory investigations. We also wish to acknowledge the Pandrillus Foundation and CERCOPAN Nigeria for giving us access to their wildlifesanctuaries during data collection.

