## Supplemental file for "Distribution of members of the *Anopheles gambiae* complex in selected forested tourist areas of Cross River State, Nigeria"

**Supplementary Table 1: Distribution of *Anopheles species* in forested area and border communities of Cross River State by multiplex PCR**.

| | Study area | No.(%) Unknown *Anopheles gambiae* complex | No.(%) *Anopheles gambiaes.s* | *No.(%) *Anopheles rufipes* | No.(%) Positive for sporozoites |
|---|---|---|---|---|---|
| Forested area | CRNP (Forest) | 3(2.9) | 31 (21.4) | 0 | 0 |
| Forested area | Rhoko Forest | 2(1.9) | 0 | 0 | 0 |
| Forested area | Drill Ranch /Afi (forest) | - | 3(1.9) | 0 | 0 |
| Border community | Obung&Aking | 18 (17.5) | 34 (23.3) | *3 (2.9) | 0 |
| Border community | Bonchor | - | 8 (7.8) | 0 | 0 |
| Wildlife Sanctuary located in an urban area | Calabar | 0 | 2(1.9) | 0 | 0 |
| Border community | Iko-esai | - | 0 | 0 | 0 |
| | Total-104(100%) | 23(22.1) | 78(75) | *3(2.9) | 0 |

***A. rufipes was identified by morphological identification keys**

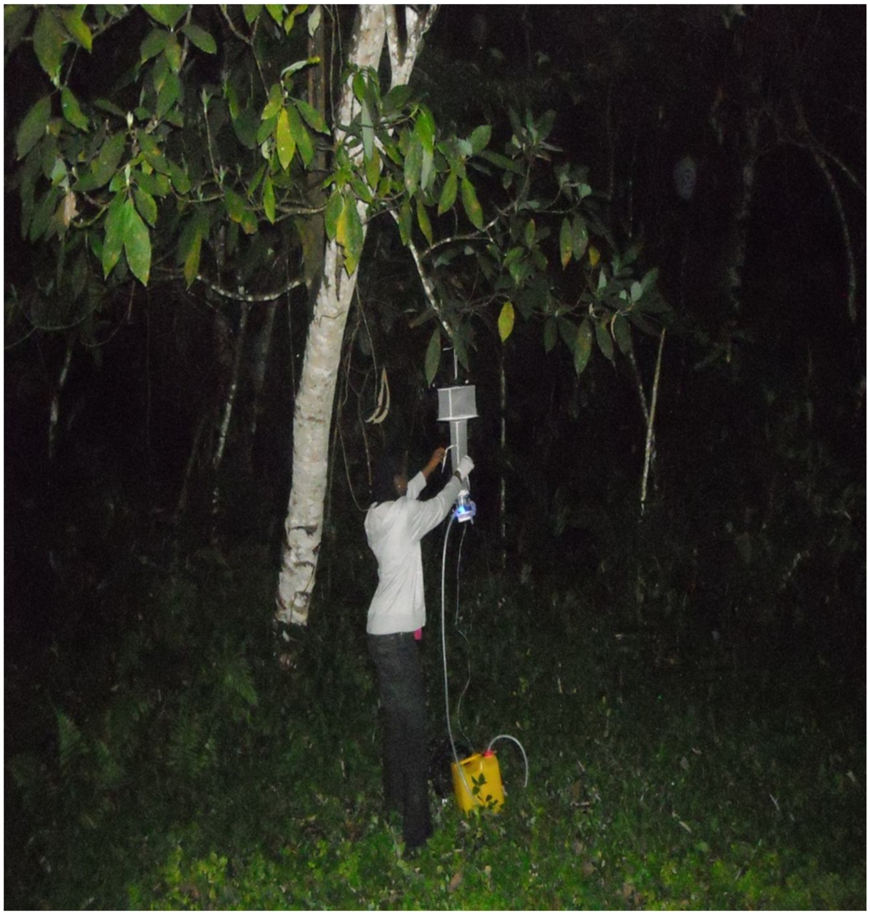

**Supplementary Figure 1: Setting up Modified CDC UV light trap at Cross River National Park**

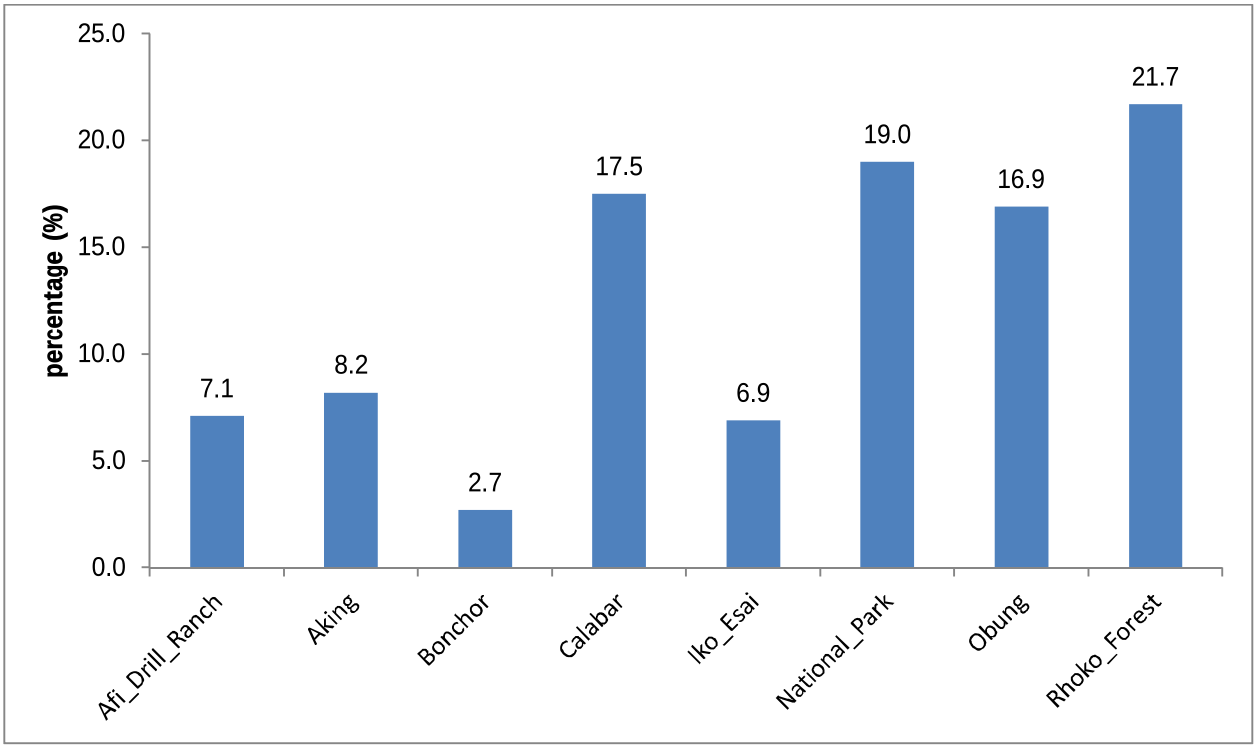


**Supplementary Figure 2 Proportion of female mosquitoes collected per study site**
